# Non-ohmic tissue conduction in cardiac electrophysiology: upscaling the non-linear voltage-dependent conductance of gap junctions

**DOI:** 10.1101/690255

**Authors:** Daniel E. Hurtado, Javiera Jilberto, Grigory Panasenko

## Abstract

Gap junctions are key mediators of the intercellular communication in cardiac tissue, and their function is vital to sustain normal cardiac electrical activity. Conduction through gap junctions strongly depends on the hemichannel arrangement and transjunctional voltage, rendering the intercellular conductance highly non-Ohmic. Despite this marked non-linear behavior, current tissue-level models of cardiac conduction are rooted on the assumption that gap-junctions conductance is constant (Ohmic), which results in inaccurate predictions of electrical propagation, particularly in the low junctional-coupling regime observed under pathological conditions. In this work, we present a novel non-Ohmic multiscale (NOM) model of cardiac conduction that is suitable for tissue-level simulations. Using non-linear homogenization theory, we develop a conductivity model that seamlessly upscales the voltage-dependent conductance of gap junctions, without the need of explicitly modeling gap junctions. The NOM model allows for the simulation of electrical propagation in tissue-level cardiac domains that accurately resemble that of cell-based microscopic models for a wide range of junctional coupling scenarios, recovering key conduction features at a fraction of the computational complexity. A unique feature of the NOM model is the possibility of upscaling the response of non-symmetric gap-junction conductance distributions, which result in conduction velocities that strongly depend on the direction of propagation, thus allowing to model the normal and retrograde conduction observed in certain regions of the heart. We envision that the NOM model will enable organ-level simulations that are informed by sub- and inter-cellular mechanisms, delivering an accurate and predictive in-silico tool for understanding the heart function.

**Author summary:** The heart relies on the propagation of electrical impulses that are mediated gap junctions, whose conduction properties vary depending on the transjunctional voltage. Despite this non-linear feature, current mathematical models assume that cardiac tissue behaves like an Ohmic (linear) material, thus delivering inaccurate results when simulated in a computer. Here we present a novel mathematical multiscale model that explicitly includes the non-Ohmic response of gap junctions in its predictions. Our results show that the proposed model recovers important conduction features modulated by gap junctions at a fraction of the computational complexity. This contribution represents an important step towards constructing computer models of a whole heart that can predict organ-level behavior in reasonable computing times.

## Introduction

The conduction of electrical waves in cardiac tissue is key to human life, as the synchronized contraction of the cardiac muscle is controlled by electrical impulses that travel in a coordinated manner throughout the heart chambers. Under pathological conditions cardiac conduction can be severely reduced, potentially leading to reentrant arrhythmias and ultimately death if normal propagation is not restored properly [1]. At a subcellular level, electrical communication in cardiac tissue occurs by means of a rapid flow of ions moving through the cytoplasm of cardiac cells, and a slower intercellular flow mediated by gap junctions embedded in the intercalated discs. Gap junctions are intercellular channels composed by hemichannels of specialized proteins, known as connexins, that control the passage of ions between neighboring cells [2]. The regulation of ionic flow through gap junctions has been established for a variety of connexin types and hexameric arrangements, which under dynamic conditions result in a markedly non-linear relation between the electric conductance and the transjunctional voltage [3], revealing a non-ohmic electrical behavior. Further, it has been shown that ionic flow through cell junctions can take up to 50% of the total conduction time in cultured strands of myocytes with normal coupling levels [4], and that conduction velocity is predominantly controlled by the level of gap-junctional communication [5], which highlights the key physiological relevance of gap-junction conductivity and coupling in tissue electrical conduction.

Cardiac modeling and simulation has strongly motivated the development of tissue-level mathematical models of electrophysiology, as they have the ability to connect subcellular mechanisms to whole-organ behavior [6]. To date, the vast majority of continuum models assume a linear conduction model of spatial communication, based on the assumption that electrical current in cardiac tissue follows Ohm’s law, i.e, that current is linearly proportional to gradients in the intracellular potential [7, 8]. From a mathematical perspective, the assumption that conduction in cardiac tissue follows Ohm’s law is conveniently represented by a linear diffusion term when stating the local statement of current balance in a continuum, where gradients are modulated by a conductivity tensor that is independent of the local electrical activity. Further, if the conductivity tensor is assumed isotropic, a Laplacian operator acting on the transmembrane potential arises, and the electrophysiology model takes the form of a set of nonlinear reaction-diffusion partial differential equations, otherwise known as the cable (monodomain) model of cardiac electrophysiology [9].

Using two-scale asymptotic homogenization techniques, analytic expressions have been obtained for the effective conductivity tensor, which is then used to model the electrical current in an average macroscopic sense [10–12]. To this end, periodicity at the microstructural level of cardiac tissue is assumed, and a representative tissue unit is partitioned in regions of high and low conductivity that represent the cytoplasm and intercalated discs with gap junctions, respectively. While this approach allows for the explicit consideration of regions with decreased conductivity, e.g. membranes where flow is mediated by gap junctions, Ohm’s law is still assumed to hold throughout the microstructural domain [13]. As a result, the non-Ohmic behavior of gap junctions and their impact on tissue-level conduction continues to be neglected [14]. In particular, it has been shown that continuum models that consider effective conductivity tensors described above fail to capture the slow conduction of electrical impulses in cases of low gap-junctional coupling [12, 15], limiting their applicability to the simulation of pathological conditions in excitable tissue. Alternatively, non-linear diffusion models that replace the laplacian term in the monodomain equations by either a fractional laplacian [16, 17] or a porous-medium-like diffusive term [8, 18], which have shown to modulate the shape of propagating waves and other restitution properties. However, these models are largely based on phenomenological grounds, and are not able to directly incorporate physical microscopic information, neither have been assessed for cases of low junctional coupling.

In this work, we present a multiscale continuum model of cardiac conduction that accounts for the nonlinear communication between adjacent cells. We argue that the explicit consideration of the non-ohmic behavior of gap junctions can be seamlessly embedded into continuum tissue-scale models of electrophysiology using an asymptotic homogenization approach, which delivers nonlinear continuum equations for characterizing the electrical conduction in excitable media.

## Results

The effect of gap-junctional coupling (GJc) on conduction is studied by down-scaling the maximal gap-junction conductance from 100% to 0.5%. The cardiac strand is excited on one end with a current whose amplitudes that varied between 10 *µ*A/mm^2^ to 35 *µ*A/mm^2^, which elicits a propagating pulse from left to right. Our results are compared with those obtained from the cell-chain model of Kucera et al. [5], which we consider as the baseline, an with the predictions from a linear homogenization model (LHM), which considers a standard cable model where the effective conductivity results from linear homogenization theory [12]. We note here that the cell-chain model results in a dynamical system with 642 degrees of freedom, using cell segments of 10 *µ*m, whereas the LHM and the HOM models employ only 65 degrees of freedom, equivalent to a spatial discretization of 100 *µ*m.

Figure 1 shows the propagating wavefronts that result from the three conduction models under different coupling levels. For GJc=100% we observe that the three models predict a very similar wavefront and wave speed. When GJc was reduced to 10% the LHM results in propagating waves that considerably drift ahead of the baseline, whereas the NOM delivers a very good prediction of the wavefront when compared to the baseline. For the case of very low coupling (GJc=1%), the NOM wavefronts display a positive drift from the baseline, but still deliver a much better prediction than the LHM model. The conduction velocity as a function of the reduction in GJc is reported in Figure 2(a). Both the LHM and NOM capture a marked decrease in conduction as the GJc is decreased, but the LHM consistently overestimates the conduction velocity, resulting in larger relative errors when compared to the NOM, particularly for GJc *<* 50%, see Figure 2(b).

**Fig 1.**
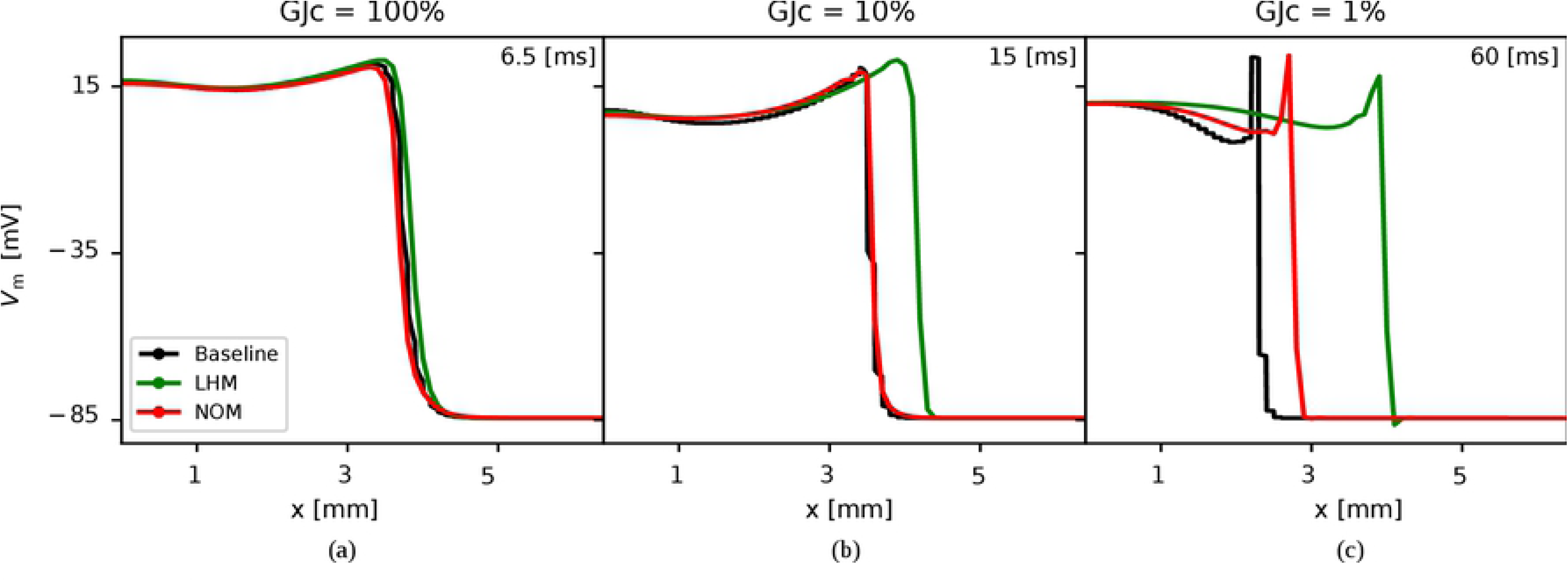
Impulse conduction features from computational simulations. The wavefront predicted by the cell-network (Baseline), linear homogenization model (LHM) and non-ohmic multiscale model (NOM) are compared for three levels of transjunctional coupling: (a) high coupling 100%, (b) low coupling 10%, and (c) very low couping 1%. Waveform and conduction speed are recovered by both LHM and HOM models at high levels of transjunctional coupling, but substantial differences can be observed at very low levels of junctional coupling, with the NOM delivering a considerably better estimate than the LHM.

**Fig 2.**
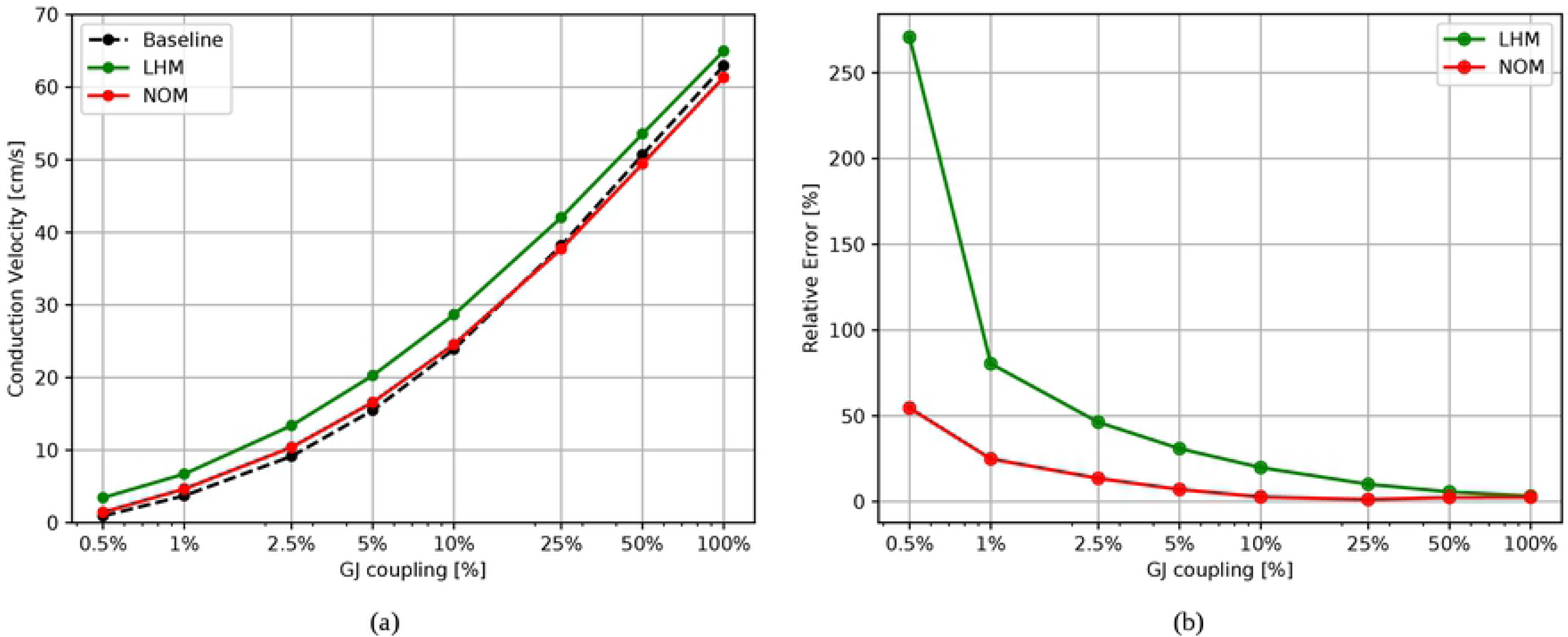
Conduction velocity studies and the effect of gap-junction coupling. (a) Performance of the LHM and the NOM as a function of the level of gap-juntional coupling, where the baseline estimates correspond to a cell-network model [5]. (b) Relative errors in conduction velocity for the LHM and NOM. At very low gap-junction coupling levels, the NOM outperforms the LHM.

The NOM model is then used to study how different type of connexins and hemichannel combinations affect the tissue-level conduction. To this end, we consider gap junctions formed by homomeric-homotypic channels Cx43-Cx43 and Cx45-Cx45, and the homomeric-heterotypic channel Cx43-Cx45, whose normalized conductance distributions are depicted in Figure 3, and whose parameters for the Boltzmann model described in (11) have been reported in the literature [3], and are summarized in Table 1. For all three types of channels, a left-to-right propagating wave is elicited by stimulating the left end of the cardiac strand. To study the effect of asymmetric conductance in reverse conduction, additional right-to-left waves are propagated for all channels studied.

**Table 1.** Parameters for the conductance distribution of gap junctions, taken from [3]. For *V*_*j*0_, *g*_*j,min*_, *z* the negative/positive values are presented. The Cx43-Cx45 case considered a modified Boltzmann distribution to improve the fitness to data.

| Channel Type | $V_{j0}$ | $g_{j,min}$ | $z$ | $p$ | $d$ |
| --- | --- | --- | --- | --- | --- |
| Cx43-Cx43 | -60.8/62.9 | 0.26/0.25 | -3.4/2.9 | 1 | 0 |
| Cx45-Cx45 | -38.9/38.5 | 0.16/0.17 | -2.5/2.7 | 1 | 0 |
| Cx43-Cx45 | -15.9/149.3 | 0.05/0.05 | -2.1/0.7 | 0.73 | 25 |

**Fig 3.**
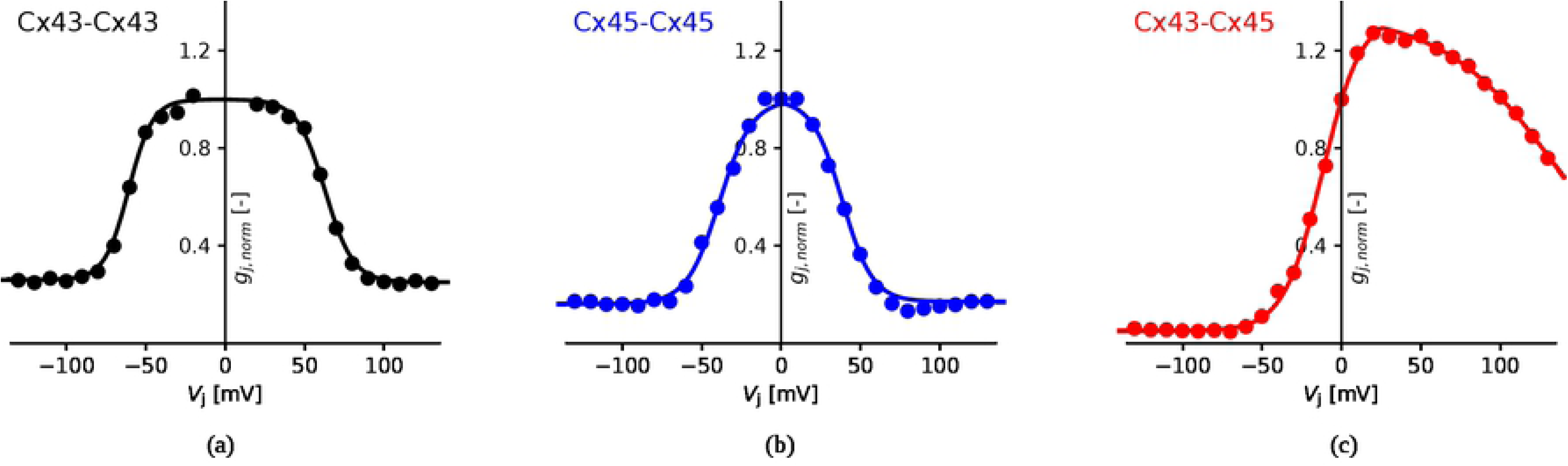
Normalized conductance of gap junctions as a function of the transjunctional voltage. (a) Cx43-Cx43 channel, (b) Cx45-Cx45 channel, and (c) Cx43-Cx45 channel. Data extracted from [3].

Figure 4(a) shows the time evolution of the transmembrane voltage measured at the middle of the cardiac strand. Activation times for the different gap-junction channels markedly differ, with Cx43-Cx43 resulting in the earliest activation, and Cx43-Cx45 in the latest activation. The conduction velocity is 59.7, 51.3 and 32.1 cm/s for the Cx43-Cx43, Cx45-Cx45 and Cx43-Cx45 channels, respectively. The time evolution for transmembrane voltage at the same location for the case of pulses traveling in opposite directions using channel Cx43-Cx45 is reported in Figure 4(b). The right-to-left wave activates this location at *t* = 6.84 ms, whereas the left-to-right wave excites the same point at *t* = 10.51 ms, which corresponds to conduction velocities of 32.2 and 63.4 cm/s, respectively. No differences in activation during reverse conduction are found for the case of homomeric-homotopyc channels Cx43-Cx43 and Cx45-Cx45.

**Fig 4.**
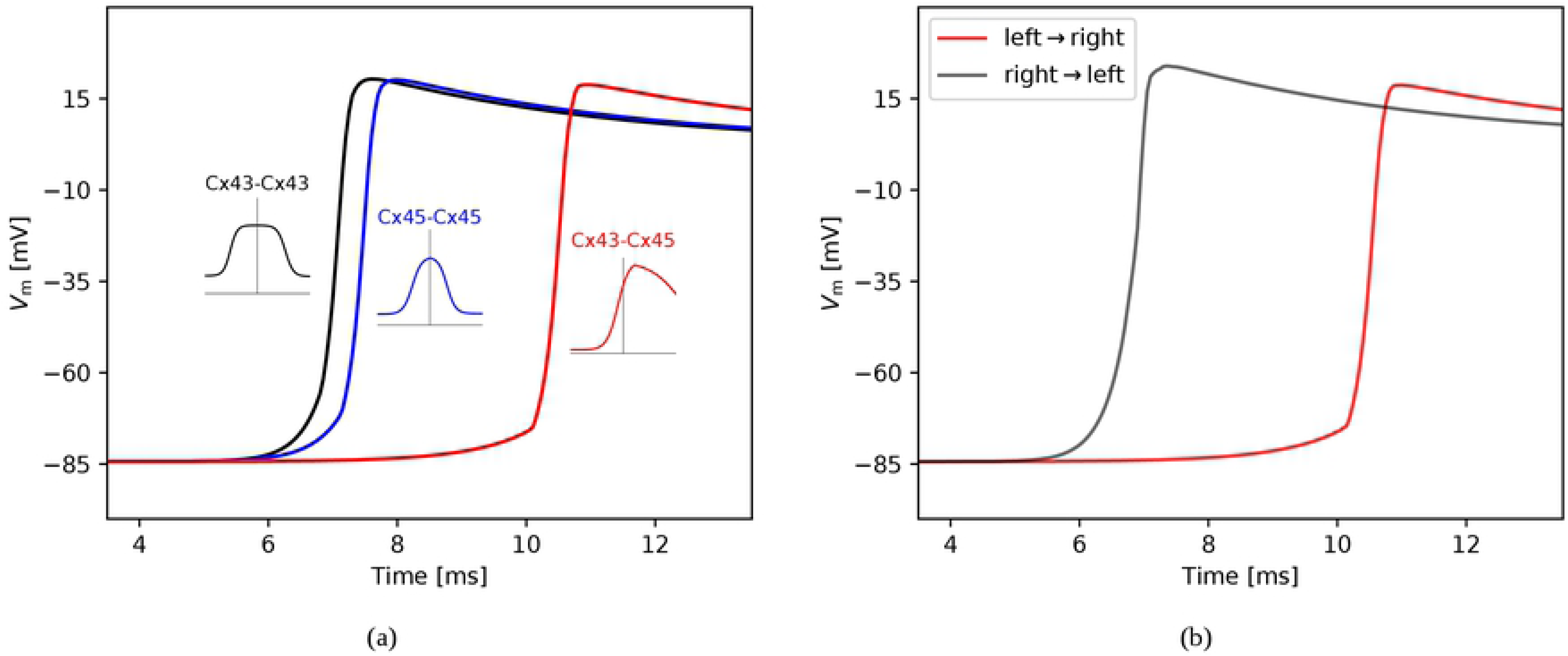
Upscaling the effect of gap-junction conductance distribution on tissue-level conduction properties. (a) Transmembrane time evolution at the center of a cardiac strand for traveling waves considering different homotypic and heterotypic channels, (b) Transmembrane time evolution for normal (left-to-right) and retrograde (right-to-left) propagation. The shape of the normalized conductance distribution results in drastic changes in the activation times. In particular, asymmetric conductance distribution predicts different conduction velocities for normal and retrograde propagation.

## Discussion

In this article, we study the gap-junction-mediated electrical conduction in excitable cardiac tissue by means of a novel non-ohmic multiscale model. A unique feature of the proposed model is that tissue-level spatial conduction is fully informed by sub-cellular communication mechanisms, specifically by cytoplasmic and gap-junctional conductances. While the upscaling of conduction properties in excitable media has been the subject of some studies in the past using a linear homogenization theory approach [10, 11], our work offers a rigorous mathematical framework that delivers an effective non-linear model of conduction able to represent, at the tissue level, the non-Ohmic conduction that takes place at the sub-cellular level. Despite the fact that our focus has been on understanding gap-mediated communication between cardiac myocytes, the present model of conduction can be extended to study the electrical propagation phenomena in other areas of biology, such as the neurosciences, where electrical synapsis occurring in the brain is highly regulated by neural gap junctions [19].

The propagating waves resulting from the NOM model resemble the wave forms and wave speeds observed in simulations of cell-strand models [5] (Figure 1). In particular, features that arise in propagating action potentials under decreasing levels of coupling such as a steeper upstroke and a notch in the upstroke [20] are predicted by the NOM model. Remarkably, this prediction is achieved at a fraction of the computational complexity involved in cell-network models, as the number of degrees of freedom in the NOM model are one order of magnitude smaller. An alternative approach is the use of hybrid multiscale models [21], which adaptively partition the domain to solve macroscopic cable equations in regions with low potential gradients and impose microscopic equations of conduction in regions of high potential gradients. While hybrid models can reduce the computational complexity of a simulation, they involve an important increase of degrees of freedom when compared to standard homogenized models. We believe that the NOM model offers the advantage of delivering accurate predictions while maintaining the computational cost similar to that of standard macroscopic continuum models. The balance between predictive power and computational cost remains one of the main hurdles in the development of patient-specific whole-heart simulations [22], which highlights the importance of developing accurate yet efficient tissue-level models.

The accuracy of the NOM and LHM models is studied by determining the conduction velocity for a wide range of gap-junctional coupling levels and comparing these results with cell-chain simulations (Figure 2). Previous studies have confirmed that the accuracy of LHM in predicting cardiac conduction consistently deteriorates as the junctional conductance is decreased to low levels [12, 15]. Remarkably, the NOM model is able to capture conduction under very low junctional-coupling scenarios with a reasonable error (Figure 2). This feature takes particular relevance in the study of cardiac disease, as the reduction of gap-junctional coupling has been correlated to a marked decreased of conduction velocity [23], and slow conduction is considered one of the main mechanisms of sustained reentrant arrhythmias [1, 24].

A unique feature of the NOM model is its ability to predict the tissue-level conduction mediated by homotypic and heterotypic combinations of homomeric connexons. In our simulations, action potentials resulting from gap-junction channels composed by homotypic Cx43-Cx43 result in a higher conduction velocity when compared to simulations considering homotypic Cx45-Cx45 channels (Figure 4(a)). This result is consistent with observations from dual whole-cell patch clamp experiments, where the conductance of Cx43-Cx43 channels can be twice as large as that of Cx45-Cx45 channels [25]. It is important to note, however, that future developments should focus in combining the gap-junction conductance distributions, as several connexin types are typically co-expressed in cardiac tissue. Another interesting feature of the NOM model is the possibility to upscale the effect of asymmetric distributions of gap-junction conductance associated to heterotypic channels (Figure 3). Here we show that such asymmetry results in propagating action potentials whose wave form and wave speed strongly depend on the direction of propagation (Figure 4(b)). This behavior, together with connexin coexpression, may partly explain the differences in conduction velocity for normal and retrograde conduction that have been observed in the sinoatrial node [26].

The work presented here can be extended in several directions. First, the theoretical framework for the NOM model should be extended to consider the 3D case of cardiac conduction. A heuristic derivation is included in Remark 2 of the S1 Appendix. Second, intercellular communication mechanisms other than gap junctions should be integrated to this theoretical framework. Sodium channels have been reported to co-localize with gap junctions at the intercalated discs, creating an ephatic coupling effect that has been associated to conduction during gap-junction blockage [24]. Further, the spatial distribution of sodium channels around the cellular membrane and on the intercalated discs has been studied using detailed cell-to-cell computational studies, to conclude that channel spatial distribution strongly affect the cardiac conduction [27]. Since the ephatic effect has been considered in homogenization schemes of cardiac conduction in the past by including a cleft-to-ground resistance in the microscopic model of conduction [12, 21], we forsee that future versions of the NOM could equally incorporate this effect, potentially in 3D formulations with non-uniform distributions of channels. Finally, the applicability of the NOM model should tested in the simulation of conduction in the whole heart during diseased conditions [22].

## Methods

### Multiscale model for non-ohmic conduction

In the following we consider the microscopic problem of non-linear conduction in a strand of cardiac cells with domain Ω = (0, *L*), see Figure 5. We let *ε* be the cell length, *δε* be the length of gap junctions, and assume that *δε* ≪ *ε* ≪ *L*. Further, we let *u*_*ε,δ*_ be the microscopic transmembrane potential field, and *j*_*ε,δ*_ be the microscopic current density. The steady-state problem of conduction resulting from current balance reads

**Fig 5.**
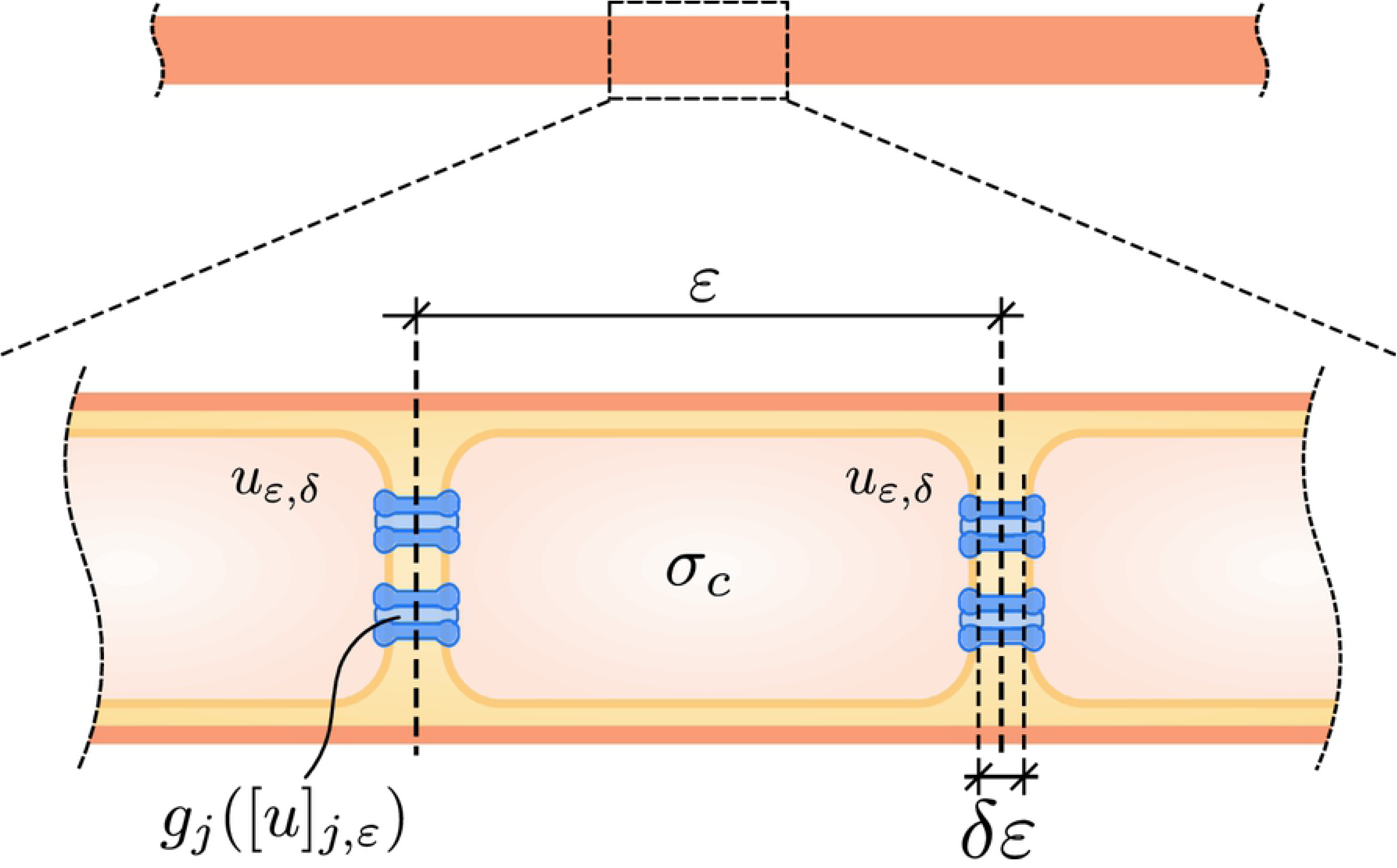
Schematic of the multiscale model of cardiac conduction. Ionic currents are linearly proportional to gradients of transmembrane potential inside the cytoplasm, but are non-linearly mediated by gap junctions located at the intercalated discs.

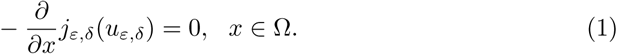

We denote the space occupied by the cytoplasm by

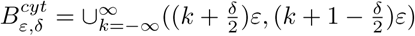, and the space occupied by gap junctions by 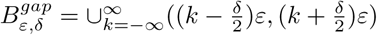. Further, we assume that current is governed by Ohm’s law inside the cytoplasm with conductivity *σ*_*c*_, but is non-linearly regulated at the gap junctions, which we express by the following microscopic constitutive law

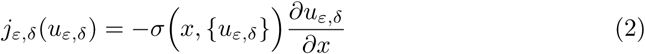

where the conductivity is described by the following relation

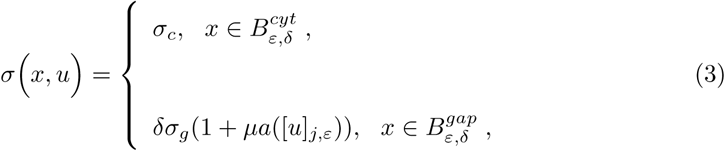

where *δσ*_*g*_ is a representative conductivity for the intercalated disc with gap junctions, *µ* is a positive constant, and *a* is a smooth bounded function that depends on the transjunctional voltage jump defined as

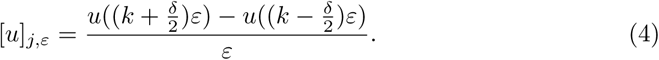

Using asymptotic analysis (see S1 Appendix for details and proofs) we show that the macroscopic current conservation for the steady-state problem is governed by the homogenized equation

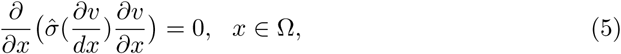

where *v* is the macroscopic transmembrane potential, and the effective conductivity modulating conduction at the macroscopic scale takes the form

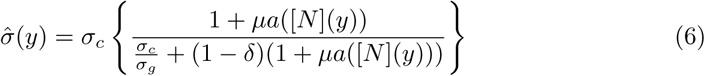

where

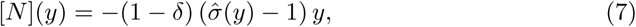

and we note that for a given transmembrane potential gradient *y*, the effective conductivity 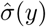 is implicitly solved from (6) and (7). Further, we show that under reasonable assumptions, the following error estimate for the macroscopic transmembrane potential holds

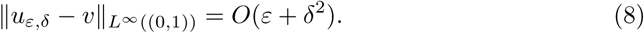

We now focus on the time-dependent macroscopic model of cardiac electrophysiology for the time interval (0, *T*). The homogenized electrical flux described in the right-hand side of (5) is then balanced by the transmembrane current, leading to the non-Ohmic cable equation

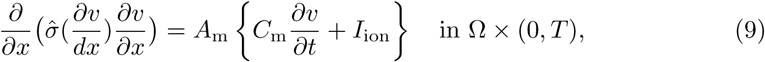

where *I*_ion_ : ℝ × ℝ → ℝ represents the transmembrane ionic current, *C*_m_ is the membrane capacity and *A*_m_ is the surface-to-volume ratio, and we note that the right-hand side of (9) accounts for the amount of charge that leaves the intracellular domain and enters the extracellular domain. Further, we will assume that the transmembrane ionic current *I*_ion_ is governed by *v* and by gating variables ***w*** : Ω × (0, *T*) → ℝ^*M*^ that modulate the conductance of ion channels, pumps and exchangers, i.e., *I*_ion_ = *I*_ion_(*v*, ***w***), where the exact functional form of *I*_ion_ will depend on the choice of ionic model. The evolution of gating variables is determined by kinetic equations of the form

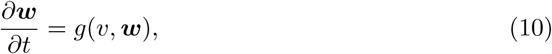

where the form of *g* : ℝ × ℝ^*M*^ → ℝ^*M*^ will also depend on chosen the ionic model. The equations (9) and (10) are supplemented with initial and boundary conditions for the transmembrane potential and gating variables to form an initial boundary value problem. The numerical solution of the coupled system of the non-Ohmic cable equation (9) and kinetic equations (10) was performed using a standard Galerkin finite-element scheme [28] for the spatial discretization and a Forward Euler scheme for the time discretization implemented in FEniCS [29], see S1 Appendix. Codes are available for download at https://github.com/dehurtado/NonOhmicConduction.

### Conduction experiments in a cardiac strand

To validate the proposed NOM model we consider the cardiac conduction problem described in [5], in which a strand of cardiac cells electrically connected by gap junctions are represented using a circuit network. The effect of gap junctions on the overall conduction is studied by varying the level of transjunctional coupling. In this work, we model the propagation of electrical impulses based on (9) and (10) in a strand with length *L* = 6.4 mm, and consider the Luo-Rudy I model of transmembrane ionic current [30]. The cytoplasmic conductivity and the membrane capacitance are taken to be *σ*_*c*_ = 0.667 *µ*S and *C*_m_ = 1 *µ*F/cm^2^ respectively [5]. The surface to volume ratio is given by *A*_m_ = 2RCG*/a*, where RCG = 2 is the ratio between capacitive and geometrical areas and *a* = 11 *µ*m is the fiber radius [5, 31]. The microscopic nonlinear normalized conductance of gap junctions is assumed to follow a Boltzmann distribution that depends on the transjunctional voltage *V*_j_, which takes the form [32]

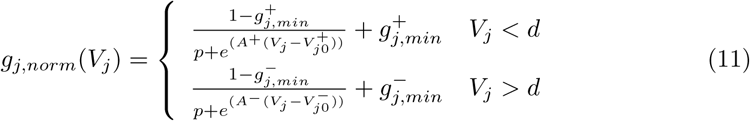

where 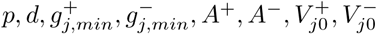 are model constants that depend on the type of channel. In this example, we consider the values reported in [3] for gap junctions based on Cx43-Cx43 channels. From (3) and the relation between electrical conductance and conductivity for a cylindrical domain we have

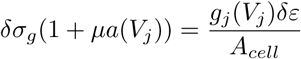

where we set *ε* = 100 *µ*m as the length of a cardiomyocyte, *δ* = 10^−4^ as the ratio between the gap length and the cell length and *A*_*cell*_ = 380 *µ*m as the transversal area of the cell. We let 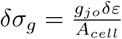, where *g*_*jo*_ is a representative conductance for the intercalated disc with gap junctions, whose value is taken to be *g*_*jo*_ = 2.534 *µ*S according to [5]. Further, we express the intercalated-disc conductance *g*_*j*_(*V*_*j*_) as the product of the gap-junction normalized conductance *g*_*j,norm*_(*V*_*j*_) times an effective conductance density and its respective plaque area, which reads

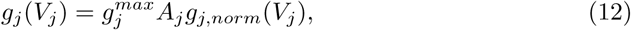

where 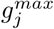 is the maximum gap-junction conductance per unit area, and *A*_*j*_ is the plaque area, i.e., the area where ions pass through gap junction channels. In our simulations, we set 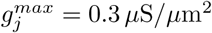 and *A*_*j*_ = 26 *µ*m^2^, both of which are within the ranges reported in the literature and provide a good fit to data [33]. As a result, we get

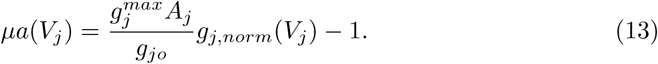

## Supporting information

## S1 Appendix. Formulation Details

Details and proofs for the asymptotic formulation and the numerical solution for the NOM model are presented here.

## Acknowledgments

The support of the Comisión Nacional de Investigación Científica y Tecnológica through their grant FONDECYT Regular #1180832 is greatly appreciated. This publication has received funding from Millenium Science Initiative of the Ministry of Economy, Development and Tourism of Chile, grant Nucleus for Cardiovascular Magnetic Resonance. The third author is supported by Russian Science Foundation grant 19-11-00033.

## Author’s contributions

D.E.H. and G.P. developed the theory, D.E.H. and J.J designed the numerical experiments, J.J. ran the simulations, D.E.H and J.J. analyzed the data, D.E.H and G.P. wrote the manuscript.

